# Affinity profiles inform clinical efficacy and tolerability of 26 antipsychotics in the treatment of schizophrenia

**DOI:** 10.64898/2026.11.19.767800

**Authors:** Benjamin Hänisch, Laila Schneider, Dirk Wildgruber

## Abstract

The pharmacological mechanisms underlying clinical efficacy and tolerability of antipsychotics in treating schizophrenia remain incompletely understood. Here, we generate multidimensional molecular affinity profiles of 26 antipsychotics or their active metabolites and relate them to meta-analytical clinical outcome data. A data-driven systematization identified two substance groups that differ significantly: The substance cluster with significantly higher affinities at 5-HT2A, 5-HT2B and 5-HT2C metabotropic serotonin receptors displayed significantly lower burden of motor-related side-effects. No single-molecule association with clinical efficacy or tolerability was significant. Canonical partial least squares analysis of clinical outcomes and affinity profiles identified two significant axes of antipsychotic diversity. One axis differentiated antipsychotics based on a motor-to-sedation side-effect continuum that separated D2 and D3 affinities mainly from affinities to metabotropic histamine and acetylcholine receptors. The second axis differentiated antipsychotics by their net treatment utility, given as a trade-off between efficacy and tolerability that cumulates in all-cause treatment discontinuation, and separated mainly metabotropic serotonin receptors affinities from affinities to metabotropic histamine and acetylcholine receptors. Together, our results highlight the importance of metabotropic serotonin antagonism for efficacious yet tolerable antipsychotic drugs.

## Introduction

Schizophrenia is a debilitating psychiatric illness and a leading global cause for psychiatric hospitalization [1,2]. Pharmacological treatment with antipsychotic drugs can effectively reduce symptoms in two thirds of patients and is an essential part of guideline-based care [3]. Despite considerable research efforts, causal links between perturbations in neurotransmitter systems and both development and treatment of schizophrenia remain elusive. Regarding pathophysiology, current explanatory hypotheses involve aberrations in multiple neurotransmitter systems, most notably dopamine [4,5], serotonin [6,7], GABA [8] and glutamate [9–11]. Similarly, it is still unclear how antipsychotic drugs alleviate symptoms of schizophrenia. While antagonism at the D2 dopamine receptor is regarded as the essential mechanism for antipsychotic effects [12], this perspective is challenged through the recent success of muscarine agonists [13] and studies showing that perturbations of other neurotransmitter systems likely play a role in mediating antipsychotic action [14,15].

The opacity of mechanisms of action contributes substantially to two interlinked major issues that complicate the clinical use of antipsychotics: First, while many different antipsychotic substances are approved for clinical use, there are no clear individual predictors for substance-dependent antipsychotic response, making large population-based studies of effectiveness an important but limited compass of drug selection. Second, and of equal importance, is the considerable burden of side-effects of antipsychotic medication, which can substantially diminish the patient’s quality of life. Side-effects are an important contributor to inadherence to prescribed medication [16], a key element of relapse and treatment failure [17]. Since differences in side-effects are generally more pronounced than differences in efficacy, side-effect profiles are the primary guideline-based substance selection mechanism [18]. Side-effects often either require symptomatic treatment, which varies in effectivity, or a switch to another antipsychotic drug, which is especially unfortunate in case of effective control of psychotic symptoms under index medication. The degree of mechanistic understanding linking side-effect profiles to pharmacodynamic characteristics is heterogeneous – while some relationships are clear, such as the link between muscarinic acetylcholine receptor blockade and anticholinergic side-effects, especially the development of motor-related symptoms like extrapyramidal symptoms or akathisia is insufficiently predictable [19,20]. Linking pharmacodynamic profiles to efficacy and tolerability is further complicated by a subset of antipsychotics that get broken down into active metabolites with sometimes relevantly different neurotransmitter receptor and transporter binding profiles, and whose role in the respective drugs’ clinical efficacy and tolerability are so far insufficiently understood.

In this study, we therefore aim to develop a clearer understanding of the relationships between the pharmacodynamic profiles of antipsychotic drugs and their clinical efficacy and tolerability. We focus on 26 antipsychotics for which multiple efficacy and tolerability measures have been assessed in a substantive meta-analysis [21]. To provide a link to their pharmacodynamic characteristics, we build multidimensional affinity profiles of antipsychotic drugs using their inhibition constants (Ki values) at various neurotransmitter transporter and receptor molecules (NTRM), which we mainly source from the PDSP Ki Database (KiDB), an open-access repository of published inhibition constants [22]. We leverage these pharmacodynamic profiles to develop a data-driven systematization of drugs, and furthermore investigate how single-molecule affinities and multivariate pharmacodynamic dimensions relate to meta-analytical clinical outcomes. To investigate the putative role of active metabolites, we conduct the main analyses using affinities of Norquetiapine and Norclozapine, active metabolites of the parent substances Quetiapine and Clozapine, which are pharmacodynamically well-characterized, but clinically poorly understood.

## Results

We first aimed to develop a data-driven systematization of antipsychotic drugs. To this end, we performed hierarchical agglomerative clustering using Ward’s linkage algorithm (**Figure 1A**) and *k*-means clustering (**Figure 1B**) of multidimensional affinity profiles in 26 efficacious substances investigated in [21]. The multidimensional affinity profiles are contained in **Supplementary Table 1**. Other than hierarchical agglomerative clustering, *k*-means clustering requires a pre-specified number of centroids. Through calculating silhouette scores across the sampled centroid space, we determined a two-cluster solution to be appropriate (**Supplementary Figure 1A**). *K*-means cluster 1 consisted of Amisulpride, Cariprazine, Haloperidol, Lurasidone, Molindone, Paliperidone, Pimozide, Sulpiride, Trifluoperazine and Thioxitene. *k*-means cluster 2 consisted of Aripirazole, Asenapine, Brexpiprazole, Chlorpromazine, Fluphenazine, Iloperazine, Loxapine, Norclozapine, Norquetiapine, Olanzapine, Perphenazine, Risperidone, Sertindole, Thioridazine, Ziprasidone and Zotepine. We repeated hierarchical agglomerative clustering using different linkage algorithms (**Supplementary Figure 1B**), where we found that the main two-cluster solution with other linkage algorithms is separating Asenapine from all other substances, making Ward’s linkage algorithm the only hierarchical agglomerative clustering method that yields an informative two-cluster solution.

**Figure 1.**
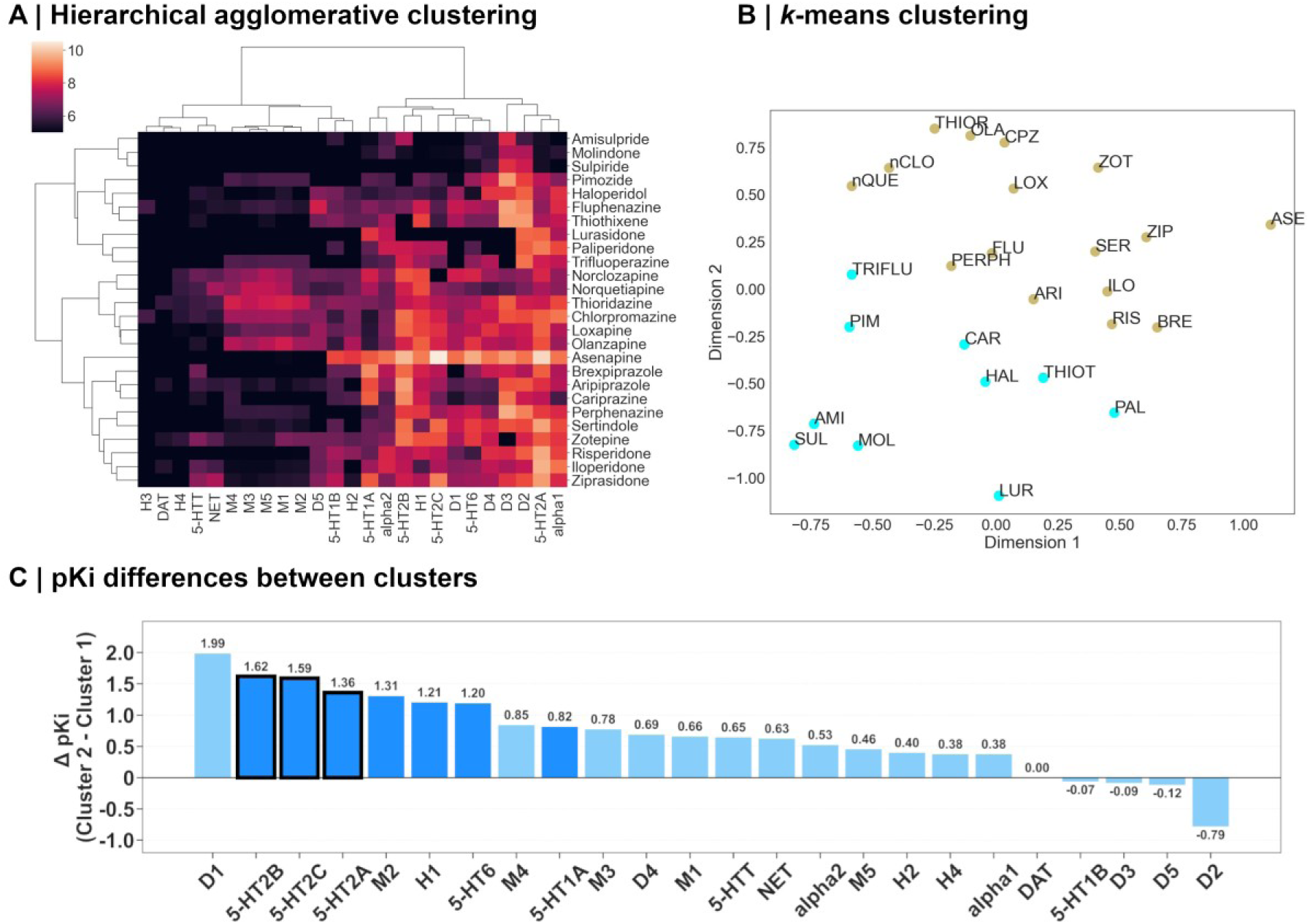
Data-driven systematization of antipsychotics based on NTRM affinity profiles. Hierarchical agglomerative clustering (A) and *k*-means clustering (B) of pKi values across 26 antipsychotics. C: Median difference in pKi values of 26 receptors between clusters. Bars in dark blue indicate nominal statistical significance. Black boxes around bars indicate False Discovery Rate (FDR)-corrected statistical significance at α=0.05 using the Benjamini-Hochberg method. AMI: Amisulpride, ARI: Aripiprazole, ASE: Asenapine, BRE: Brexpiprazole, CAR: Cariprazine, nCLO: Norclozapine, CPZ: Chlorpromazine, FLU: Fluphenazine, HAL: Haloperidol, ILO: Iloperidone, LOX: Loxapine, LUR: Lurasidone, MOL: Molindone, OLA: Olanzapine, PAL: Paliperidone, PERPH: Perphenazine, PIM: Pimozide, nQUE: Norquetiapine, RIS: Risperidone, SER: Sertindole, SUL: Sulpiride, THIOR: Thioridazine, THIOT: Thiothixene, TRIFLU: Trifluoperazine, ZIP: Ziprasidone, ZOT: Zotepine.

Group assignment between hierarchical agglomerative clustering with Ward’s linkage algorithm and *k*-means clustering largely overlapped, with the only disagreement being in the cluster assignments of Cariprazine and Fluphenazine. Following, we use hierarchical agglomerative clustering-based cluster assignments. The correspondence between the affinity-based clusters and the canonical typical/atypical distinctions as well as the more recent neuroscience-based nomenclature is displayed in **Supplementary Figure 2**.

To understand which differences in molecular profiles drive the separation into clusters, we compared median single-molecule affinities between clusters. We found False Discovery Rate (FDR)-corrected significant differences of 5-HT2B, 5-HT2C and 5-HT2A affinities between Cluster 1 and 2. Drugs in Cluster 2 had significantly *higher* median pKi values at these receptors (**Figure 1C**).

We next used this two-cluster systematization to investigate whether the clusters meaningfully differ in clinical attributes. Univariate testing between meta-analytical clinical outcome parameters showed FDR-corrected statistically significant differences in “Akathisia” and “Use of antiparkinsonian medication” between Cluster 1 and Cluster 2. Cluster 1 had significantly higher values than Cluster 2, indicating that antipsychotics in Cluster 1 cause significantly *more* “Akathisia” and “Use of antiparkinsonian medication” **(Table 1).**

**Table 1.** Clinical outcome parameter comparison between clusters. Comparison of cluster medians per outcome measure. Median burden of “Akathisia” and “Use of antiparkinsonian medication” (highlighted in bold) is significantly higher in Cluster 1, as determined using Mann Whitney U-Tests and Benjamini-Hochberg FDR correction at α = 0.05.

| Outcome | n cluster 1 | median cluster 1 | n cluster 2 | median cluster 2 | U statistic | p-value | FDR-corrected p-value |
|---|---|---|---|---|---|---|---|
| ***Akathisia*** | 10 | 4,38 | 16 | 1,99 | 139 | 0,002 | 0,026 |
| ***Use of antiparkinsonian medication*** | 10 | 3,06 | 16 | 1,565 | 134 | 0,0048 | 0,0312 |
| *At least one anticholinergic side-effect* | 10 | 1,315 | 16 | 1,695 | 47 | 0,0867 | 0,3071 |
| *Prolactin elevation in ng per ml* | 5 | 18,49 | 13 | 2,75 | 50 | 0,0945 | 0,3071 |
| *Sedation* | 10 | 1,545 | 16 | 2,17 | 55 | 0,1965 | 0,5109 |
| *Change in positive symptoms* | 4 | -0,51 | 14 | -0,415 | 17,5 | 0,2875 | 0,5986 |
| *Weight gain in kg* | 7 | 0,84 | 14 | 1,665 | 35 | 0,3223 | 0,5986 |
| *All-cause discontinuation* | 10 | 0,885 | 16 | 0,82 | 95 | 0,4443 | 0,722 |
| *QTc prolongation in ms* | 4 | 1,45 | 10 | 4,53 | 15 | 0,5395 | 0,7682 |
| *Change in depressive symptoms* | 9 | -0,2 | 14 | -0,235 | 72 | 0,5919 | 0,7682 |
| *Overall change in symptoms* | 10 | -0,445 | 16 | -0,43 | 88,5 | 0,6731 | 0,7682 |
| *Change in negative symptoms* | 4 | -0,33 | 14 | -0,36 | 32 | 0,7091 | 0,7682 |
| *Change in social functioning* | 3 | -0,44 | 8 | -0,24 | 11 | 0,9212 | 0,9212 |

We repeated the clustering procedure using the affinities of Clozapine and Quetiapine instead of the affinities of their active metabolites Norclozapine and Norquetiapine. With the exception of Quetiapine being assigned to Cluster 1 instead of Cluster 2, cluster assignments remained stable. With non-metabolised compounds, the significant clinical differences between clusters were no longer discernible, while the affinity-based differences remained stable (**Supplementary Figure 3, Supplementary Table 2**).

Following the clustering-based analysis, we investigated whether single-molecule affinities were informative regarding meta-analytical clinical outcomes through performing correlational analyses between per-substance molecular affinity profiles and clinical outcome data [21]. We excluded affinities for NET, DAT, H3 and H4 due to insufficient overlap between pKi values for these NTRM and clinical efficacy and tolerability data per drug. The original meta-analytical clinical data reports drug effects as differences from placebo, with values below a threshold indicating that the antipsychotic drug should be favoured, and values above a threshold indicating that placebo treatment should be favoured. This leads to a non-uniform directional coding of drug effects between efficacy and tolerability parameters, as drug action causing side-effects is generally coded as an above-threshold value, while drug action mediating efficacy is generally coded as a below-threshold value. As this differential coding of directionality in drug effects between efficacy and tolerability would make subsequent analyses difficult to interpret, we inverted the sign of the efficacy columns to uniformly represent a stronger drug effect as a higher value in the clinical outcome. We found various nominally statistically significant Spearman rank correlations between clinical efficacy (**Figure 2A**) or tolerability parameters (**Figure 2B**) and single-molecule pKi values, of which none survive FDR correction at α = 0.05.

**Figure 2.**
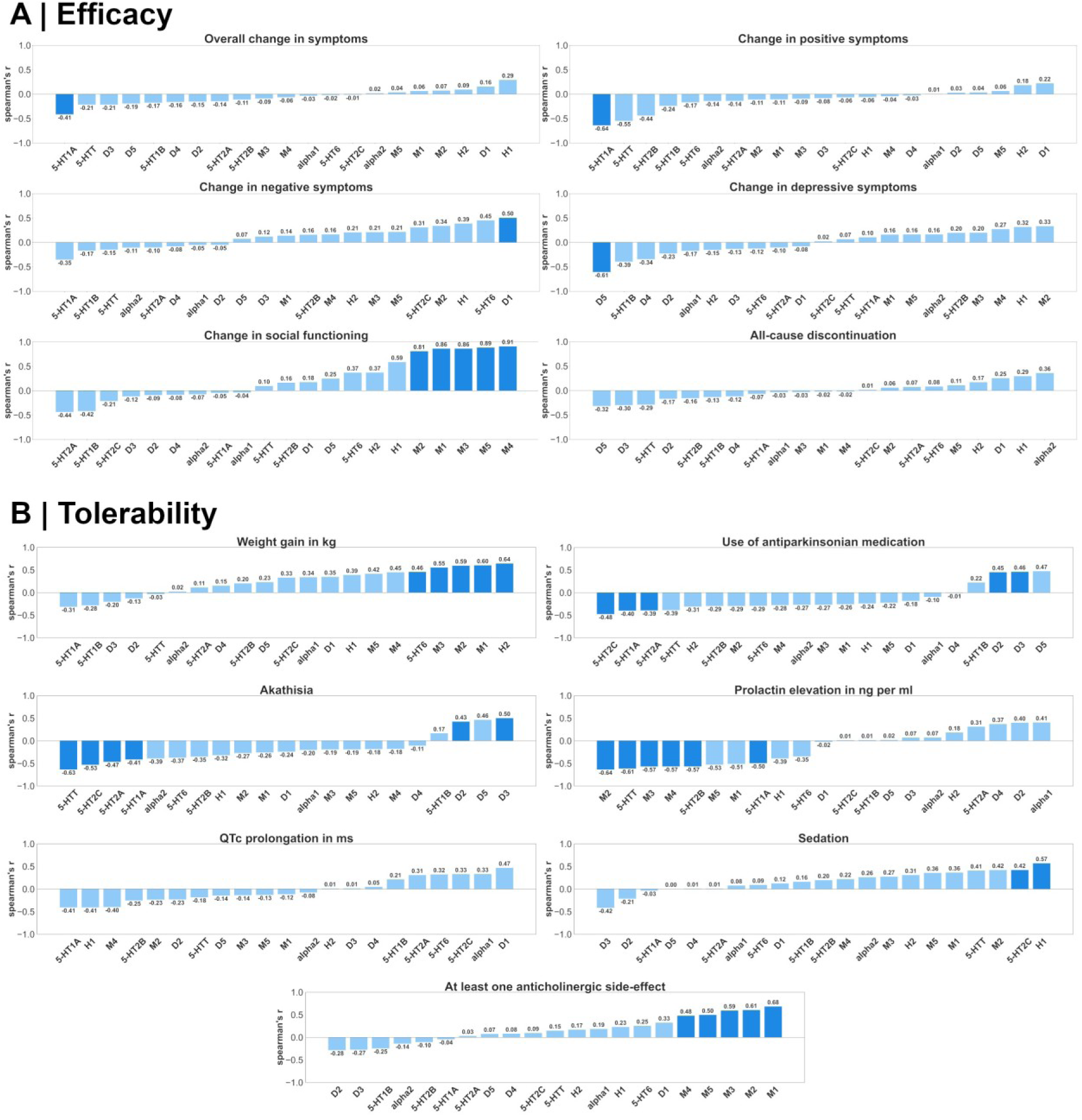
Relationship between single-molecule affinities and meta-analytical clinical outcomes. Spearman rank correlation coefficients between meta-analytical clinical efficacy (A) and tolerability (B) data and NTRM affinity profiles, expressed as pKi values, across 26 antipsychotics. Bars in dark blue indicate nominal statistical significance.

The significant differences in both clinical outcomes and metabotropic serotonin receptor affinities between clusters, combined with the absence of significant correlations between single-molecule affinities and clinical outcomes, hints at a more complex, multivariate relationship between clinical outcomes and pharmacodynamic profiles. Therefore, to further characterize the joint, multivariate structure of relationships between NTRM affinities and clinical outcomes, we performed Canonical Partial Least Squares (PLS) analysis of affinity profiles, expressed as pKi values, and meta-analytical clinical outcomes (see **Methods**). As in the single-molecule correlation analyses, we changed the sign of clinical efficacy columns so that a larger number designates a more pronounced drug effect to provide consistent coding of effect sizes and directions across outcomes. We found two statistically significant components **(Figure 3A)**. To further understand the potential clinical significance of the molecularly derived clusters, we investigated the positioning of antipsychotics in PLS-derived latent space and colour-coded them by cluster membership **(Figure 3B)**. Overall, substances that were assigned to the same cluster based solely on their molecular affinity profiles retain proximity and separability in PLS-derived latent space. Interestingly, almost all substances from Cluster 1 negatively score on Component 1, and most have a negative score on Component 2. To further understand how clinical outcomes and molecular affinities jointly drive the latent dimensions, we investigated the respective clinical and affinity loadings of the significant components **(Figure 3C)**. Component 1 differentiated “Prolactin elevation in ng per ml”, “Akathisia” and “Use of antiparkinsonian medication”, which load negatively, from the remaining meta-analytical clinical outcomes, which load positively. Affinities do D2 and D3 loaded negatively, and the remaining affinities loaded positively. Component 2 differentiated “All-cause discontinuation” and “Prolactin elevation in ng per ml”, which load positively, from the remaining meta-analytical clinical outcomes. Regarding affinities, mainly serotonin, norepinephrine and dopamine receptors and transporters – except for D3 and D5 – loaded positively, while muscarine and histamine affinities – except for H1 and H2 – loaded negatively.

**Figure 3.**
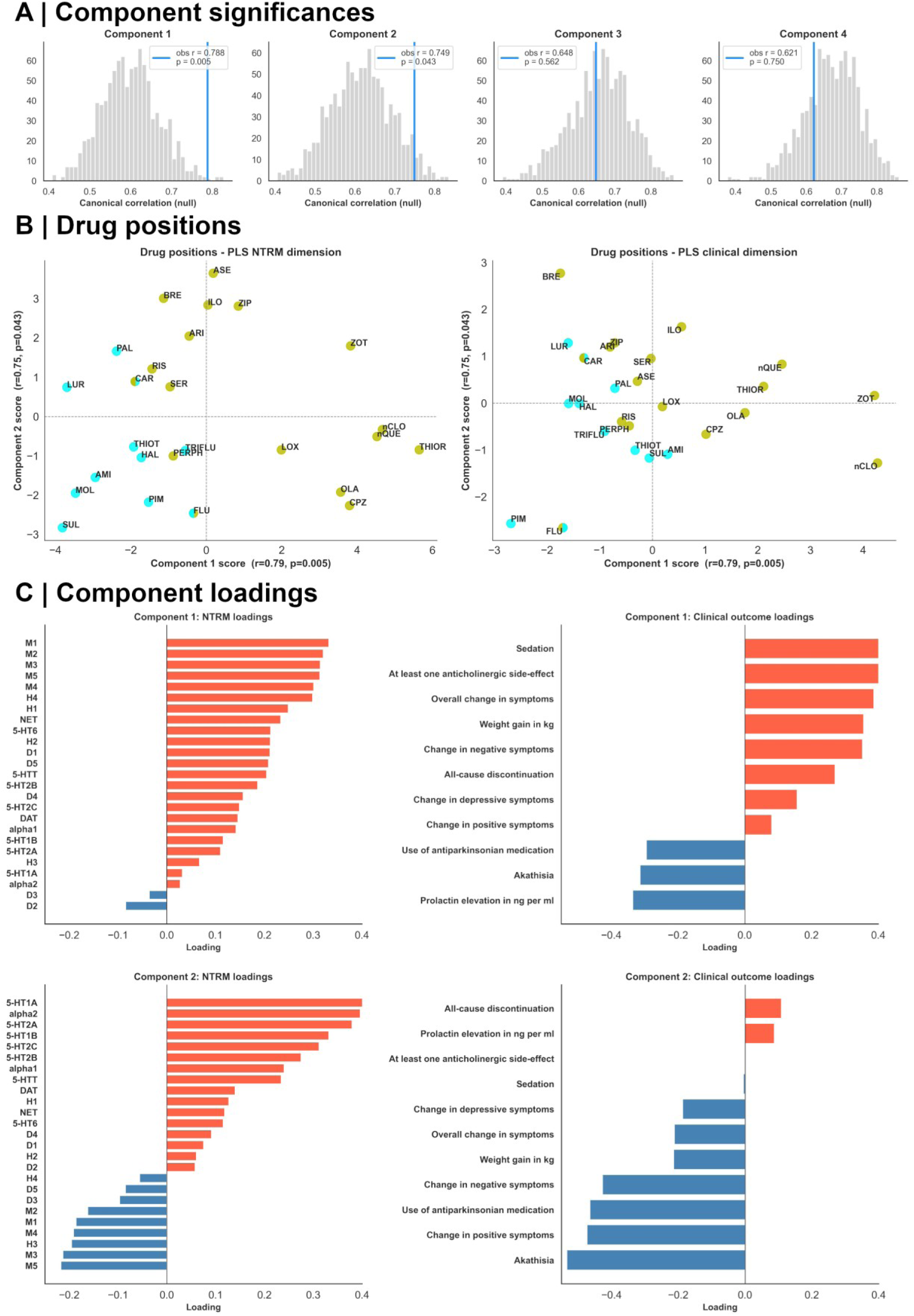
Canonical PLS analysis of NTRM affinity profiles and meta-analytical outcomes. A: Significances of canonical components. Components 1 and 2 are statistically significant as deemed by permutation testing; B: Drugs in PLS latent space as their positions on the NTRM affinity space (left) or clinical outcome space (right) on the significant components 1 and 2. Colour coding matches consensus cluster assignment. Conflicting assignment of Cariprazine and Fluphenazine is denoted by split colouring. C: Loadings of meta-analytical NTRM affinities and clinical outcomes on significant PLS components 1 and 2. Positive loadings are displayed in red, negative loadings are displayed in blue.

## Discussion

In this work, we investigated the relationship between antipsychotics’ multidimensional molecular affinity profiles and their clinical efficacy and tolerability. Using molecular affinities, we differentiated two clusters of antipsychotics that were clinically informative with regards to motor side-effects and differed significantly in median affinities to metabotropic 5-HT2A, 5-HT2B and 5-HT2C receptors. Canonical PLS analysis of NTRM affinity profiles and clinical outcomes across drugs delineated two significant components with distinct clinical and pharmacodynamic interpretability.

Our data-driven systematization of antipsychotics yielded a coarse two-group solution. Multiple taxonomies for antipsychotics exist, grouping them either by their year of introduction into first or second generation, their specificity in dopamine D2 receptor binding into typical or atypical [23], or by more specific monoamine affinities [24]. These systematization efforts reflect the hope that group or cluster membership is informative of clinical properties, which might be especially important in cases when antipsychotics need to be switched due to non-response or side-effects. Here, such systematizations may be helpful for avoiding the re-occurrence of undesired side-effects through in-group generalization of clinical properties, or to target a relevantly different neurochemical profile through between-group separation of pharmacodynamic characteristics. Other approaches have leveraged clustering approaches of affinity profiles to delineate group boundaries with some success towards achieving that aim [25], but have still relied more on expert knowledge than data-driven metrics to select the exact number of groups to form. We improved upon previous efforts in that regard through employing multiple clustering algorithms and clustering quality evaluation scores. Critically, using the affinity data of active metabolites of Clozapine and Quetiapine instead of the parent compounds, the affinity-derived groups constitute a clinically informative data-driven separation, as clusters differ significantly with regards to “Akathisia” and “Use of antiparkinsonian medication”, which can be summarised as antipsychotics-induced extrapyramidal symptoms (EPMS). Affinity enrichment per cluster sheds light at which differential affinities drive the separation and might therefore indirectly inform what drives the clinical effect. We found significantly higher affinities to the metabotropic serotonin receptors 5-HT2A, 5-HT2B and 5-HT2C. While this aligns with the canonical explanation of atypicality, which posits that concurrent 5-HT2A inhibition upon D2 blockade increases midbrain dopamine release, thereby reducing EPMS, both our clustering results as well as single-molecule affinity correlations suggest that additional mechanisms involving a broader serotonergic manipulation through 5-HT2B, 5-HT2C and potentially even 5-HTT inhibition may be involved in mediating the reduction of extrapyramidal symptoms. This is in-line with animal experiments demonstrating the effectivity of 5-HT2C antagonism in reducing antipsychotics-induced EPMS [26] and the successful treatment of EPMS through Mirtazapine, which acts at multiple metabotropic serotonin receptors [27]. Our findings may be interpreted as exploratory evidence that other drugs acting at 5-HT2B, 5-HT2C and potentially even 5-HTT may be candidate substances for the treatment of antipsychotics-associated EPMS.

Furthermore, we developed an informative multivariate perspective on the relationships between affinity profiles and clinical efficacy and tolerability through Canonical PLS analysis, which yielded two significant components with distinct and informative loadings of NTRM affinities and clinical outcomes. Clinical outcome and affinity loadings of component 1 strongly suggest to interpret it as differentiating drugs with motor-related side-effects that are associated with higher D2/D3 affinity, from drugs with sedating and anticholinergic side effects that are associated with higher H- and M-receptor affinities. This clear differentiation of side-effects and the associated differences in neurochemical profiles constitute the main axis of antipsychotic diversity and aligns with established clinical recommendations and knowledge [28]. Component 2, on the other hand, describes a more novel secondary axis of antipsychotic diversity. Using factor loadings of clinical outcome parameters and molecular affinities, it may be best interpreted as describing an axis of overall treatment utility. Critical for this interpretation is a close look at ‘All-cause discontinuation’, which summarizes discontinuation both due to treatment inefficacy (which the authors of the original meta-analysis name as the reason for discontinuation in the majority of cases, hence its treatment as an efficacy parameter) and side-effect burden. It can thus be seen as the net benefit of antipsychotic treatment and is associated with broad-spectrum affinities to dopamine, noradrenaline and serotonin receptors and transporters through their concurrent positive loading on component 2. Importantly, loadings of metabotropic serotonin receptors are more pronounced than loadings of the D2 receptor, suggesting that such an overall net benefit is primarily driven by affinities to metabotropic serotonin receptors rather than to D2. Together with our previous results, this would hint at the importance of reducing side-effects over an increased antidopaminergic potency to elevate the net benefit of antipsychotic treatment, since we find no evidence that affinity to serotonin receptors increases therapeutic efficacy. The importance of side-effect reduction to increase the overall net benefit is reinforced by the negative loadings on component 2, which group all clinical outcome parameters other than “All-cause discontinuation” and “Prolactin elevation in ng per ml” with histamine and muscarine receptor affinities. Antagonism at these receptors mediates metabolic, sedating and anticholinergic side-effects while neither reducing unwanted effects of dopamine blockade nor mediating treatment efficacy, making them weigh on the net benefit of antipsychotic treatment rather than adding to it. Finally, the negative clinical factor loadings suggest that gaining overall clinical efficacy comes at the expense of increased side-effect burden that might, in the end, not be of net benefit to the patient. The PLS coordinate space can therefore be interpreted as follows: Along component 1, antipsychotics can be selected by side-effect profile, while along component 2, antipsychotics can be selected by their overall benefit, constructed as a trade-off between side-effect burden and clinical efficacy.

Importantly, side-effects remained the primary delineating factor between affinity-based clusters and Canonical PLS components. We found no clear evidence that singular or combined molecular affinities meaningfully inform differences in clinical efficacy. Furthermore, developing clinically informative clusters was only possible upon using the affinity profiles of metabolites of Clozapine and Quetiapine. This might hint at the metabolic active compounds contributing more to the tolerability of these drugs than to their efficacy, as the metabolites are generally more serotonin-affine, which our findings suggest is more relevant for increasing tolerability than efficacy. Overall, the clear and differentiated relationship established between clinical efficacy, tolerability and molecular profiles might be useful for efforts that aim to develop novel antipsychotic drugs. Here, especially PLS component 2 could serve as an informative baseline on how molecular affinities influence treatment efficacy and tolerability trade-offs.

The results of this study might be understood as an extension of current pharmacological models, and do not touch upon more experimental strands relating neuromodulatory systems to schizophrenia and antipsychotic treatment, such as the novel approach of muscarine agonism. Further notable alternative perspectives concern glutamate-receptor focused treatment approaches. Here, especially the glutamatergic NMDA receptor has received considerable attention. Pharmacological NMDA receptor antagonism causes schizophrenia-like symptoms [11], and anti-NMDA receptor antibodies are associated with endogenous psychosis in autoimmune diseases [9,29]. Since no Ki values from human subjects for desired glutamate receptors were present in KiDB, however, investigating how direct modulation of glutamate or GABA signaling by existing antipsychotic drugs might contribute to their efficacy and tolerability was not possible in this study, and sensible interpretations how indirect influences of, for example, dopaminergic signaling might perturb these neuromodulatory systems goes beyond the scope of our methodology [30]. Once the necessary data is available, future studies can expand the current approach to also include acetylcholinergic, GABAergic and glutamatergic neuromodulatory systems.

There are several important limitations to consider when interpreting this study. First, we want to point out that KiDB-reported Ki values for the same ligand-target pair can vary considerably, which we aim to mitigate by using the median Ki value. Furthermore, the quality of studies and reporting within the database are heterogeneous, and our study inherits these issues. Additionally, the coverage of receptors and transporters varies between drugs, leading to a heterogeneous spread of missing values, which we treat as no relevant ligand-drug interaction for the clustering and Canonical PLS analyses, but which might not always reflect a true absence of interaction. Moreover, while we investigate the differential role of Norquetiapine and Norclozapine as active metabolites, several of the other investigated antipsychotics are metabolized to secondarily active compounds with sometimes relevantly different affinity profiles as well, something we cannot properly account for in this study when establishing relationships between clinical efficacy and tolerability and binding affinities. Furthermore, we want to emphasize that Ki values reflect molecular affinities and do not indicate in-vivo receptor occupancy, and molecular affinities do not allow for direct claims about drug effects realized in the brain. While the majority of reported substance-receptor bindings confer antagonistic effects, a subset of drugs and receptors, especially Aripirazole, Brexpiprazole and Cariprazine can act as partial agonists, especially at metabotropic serotonin and D2/D3 receptors [31]. Partial agonists produce sub-maximal biological responses compared to an agonist upon receptor binding. Critically, partial agonists occupy a *de facto* undefined hybrid role and act as functional *antagonists* in the presence of full agonists through receptor blocking. Whether a partial agonist realizes an agonistic or antagonistic function is thus context-dependent [32]. Therefore, while some drugs have demonstrable partial agonist activity at a subset of receptors under isolated experimental conditions, the extent to which this agonistic activity is actually expressed as a clinically relevant net agonist effect in vivo remains unclear, especially given the high clinical similarity between the aforementioned drugs with partial agonist qualities and regular receptor antagonists [21,33]. The lack of relevant difference is further strengthened by the recoverable separability of purely affinity-based clustering patterns along the Canonical PLS-derived significant components which, critically, also include relevant clinical information. We therefore chose not to account for affinities potentially realizing partial agonistic effects differently in our analyses. Given these important limitations considering the data sources and the conceptual distance between molecular affinities and in-vivo drug effect, this study is best understood as an exploratory analysis yielding informed hypotheses for potential clinical verification.

In sum, our results provide a data-driven, clinically informative systematization of antipsychotic drugs. We develop a detailed picture on the molecular correlates of antipsychotic side-effects and efficacy, highlighting the role of serotonin affinities in antipsychotic tolerability. Our study advocates for a refined perspective on atypicality – clinically, it primarily influences side-effects rather than efficacy, while on the molecular level, it should be broadened beyond 5-HT2A antagonism [34].

## Methods

### Pharmacodynamic and clinical data preparation

To generate molecular affinity profiles, we sourced inhibition constants from KiDB, an open-access repository that collects published inhibition constants [22]. Only Ki values from sources flagged as ‘human’ were used. For each substance-transporter/receptor interaction, the median of reported Ki values was calculated across studies after excluding studies that reported insignificant associations, which we defined as nanomolar Ki > 10 000, resulting in a cap of nanomolar Ki at 10 000, which was also used to represent missing data in clustering and Canonical PLS analyses. Six substances from the original meta-analysis (Clopenthixol, Flupentixol, Levomepromazine, Penfluridol, Perazine and Zuclopenthixol) were excluded due to insufficient inhibitory constants being present in the database. Affinities for Paliperidone were taken from [35], and affinities for Asenapine were extracted from the PubChem database. Affinities for Norquetiapine were sourced from [36]. The final affinity table is presented in **Supplementary Table 1.** For convenience in the analyses, we converted nanomolar Ki values to pKi values using the standard formula *pKi* = −*log*_10_(*Ki* ⋅ 10^−9^). A higher pKi value expresses greater affinity.

To assess clinical efficacy and tolerability of antipsychotics, we extracted effect sizes, which were either reported as mean difference, standardized mean difference or risk ratio in comparison to placebo, from [21], a network meta-analysis.

### Clustering

We performed clustering on molecular affinity profiles, expressed as pKi values, with both hierarchical agglomerative clustering, using Euclidean distance and various linkage algorithms, and *k*-means clustering. For *k*-means clustering, silhouette coefficients, defined as 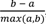, where *a* is the mean intra-cluster distance and *b* is the mean nearest-cluster distance, were used to investigate the quality of clustering solution using two to five centroids.

### Partial least squares analysis

We used Canonical Partial Least Squares analysis to examine multivariate relationships between meta-analytical clinical outcomes and multidimensional affinity profiles in antipsychotics. Canonical PLS reduces two sets of variables, subsequently referred to as X and Y, into smaller sets of orthogonal latent components that maximize the covariance between them. The NTRM affinity matrix (X) used pKi values with missing values filled to a value of 5, representing the cut-off considered as ‘no relevant affinity’. Meta-analytical clinical outcomes (Y) with more than 35% missing data across drugs were excluded, and remaining missing values were imputed with the respective outcome’s mean across substances. Before model fitting, both X and Y matrices were normalized through z-scoring.

PLS was fitted separately with three to five components. The statistical significance of each component was assessed via permutation testing (1 000 permutations): in each permutation iteration, the rows of Y were randomly shuffled to break the X–Y association, followed by model refitting. This generated a null distribution of canonical correlations per component. Subsequently, per component, we assessed the proportion of canonical correlations from the null distribution whose absolute value equaled or exceeded the observed value to compute a permutation p-value. Components with permutation p-value < 0.05 were considered statistically significant.

## Ethics declaration

This study re-analyzed previously published open-access data. No subject-level data was used. It was approved by the ethics committee of the medical faculty, University of Tübingen, Germany (Project Number 492/2024BO2).

## Acknowledgements

We thank Tobias Kaufmann and Sofie L. Valk for discussions on the subject matter. B.H. acknowledges funding from the Max Planck School of Cognition’s Clinician Scientist programm.

## Author contribution statement

BH: Conceptualization, Data curation, Formal analysis, Investigation, Methodology, Project administration, Software, Supervision, Validation, Visualization, Writing - Original Draft, Writing - Review & Editing; LS: Investigation, Writing - Review & Editing; DW: Conceptualization, Formal analysis, Investigation, Writing - Original Draft, Writing - Review & Editing;

## Code and data availability

The code and other resources to reproduce the main figure results are available at https://github.com/bhaenisch/nchem_antipsychotics_clinical

## Supplementary Figures

**Supplementary Figure 1.**
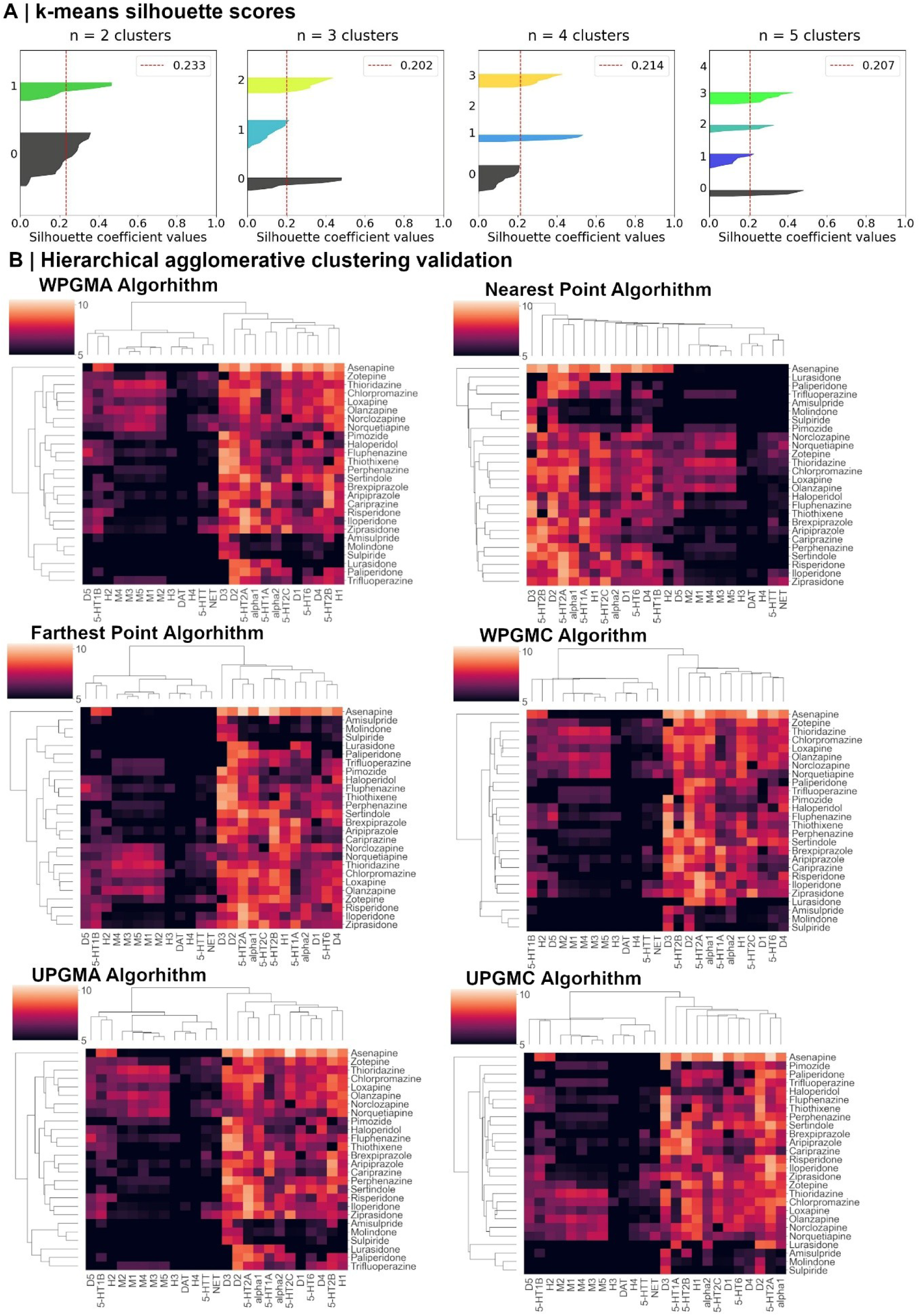
Clustering validation. A: Silhouette scores of *k*-means clustering solution using two to five centroids. B: Replication of hierarchical agglomerative clustering using different linkage algorithms.

**Supplementary Figure 2.**
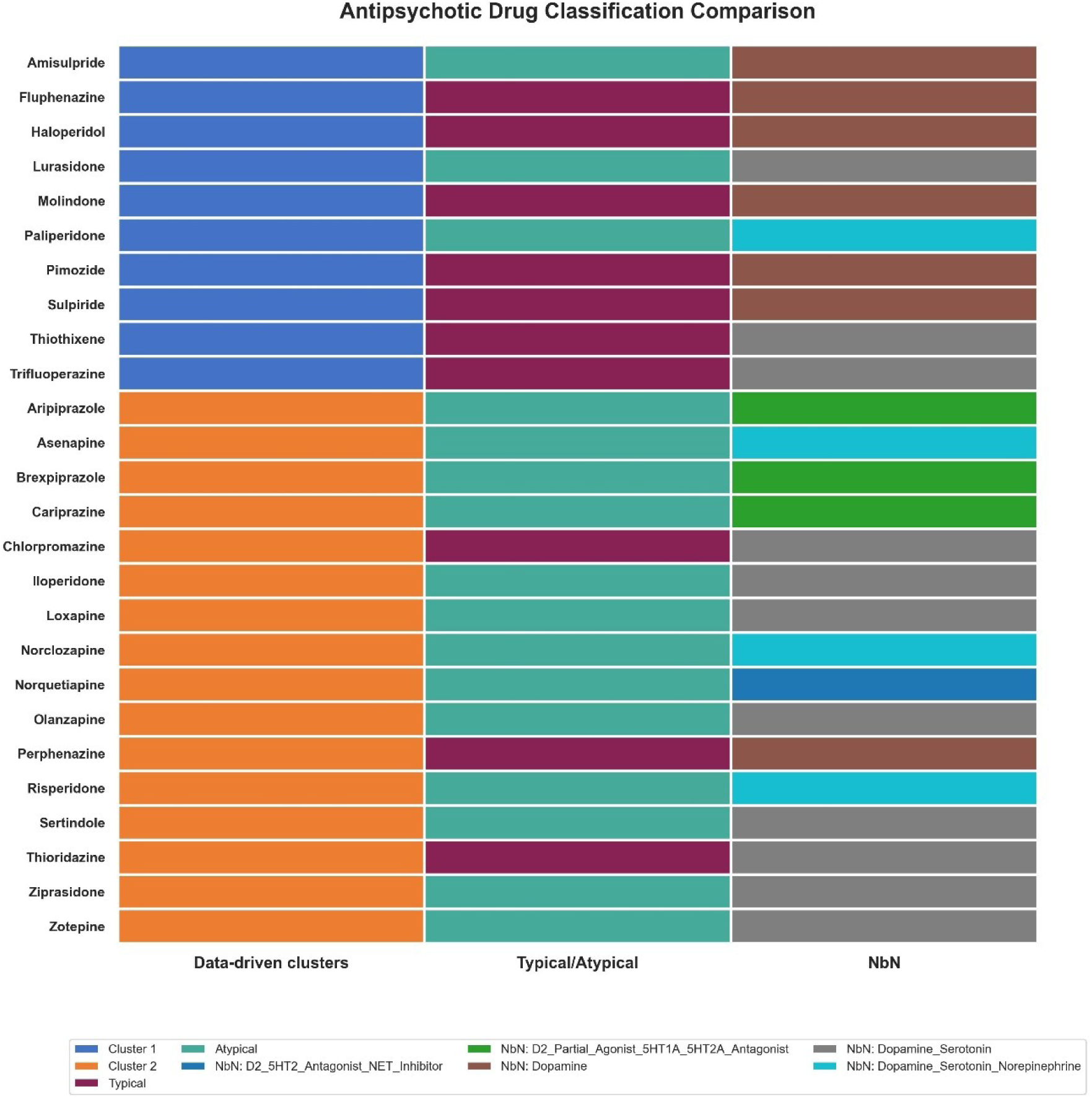
Comparison of antipsychotic groupings. The substance assignments to groups are compared between the data-driven clusters developed in this study (left), the typical/atypical distinction (middle) and neuroscience-based nomenclature (NbN; right) substance classes.

**Supplementary Figure 3.**
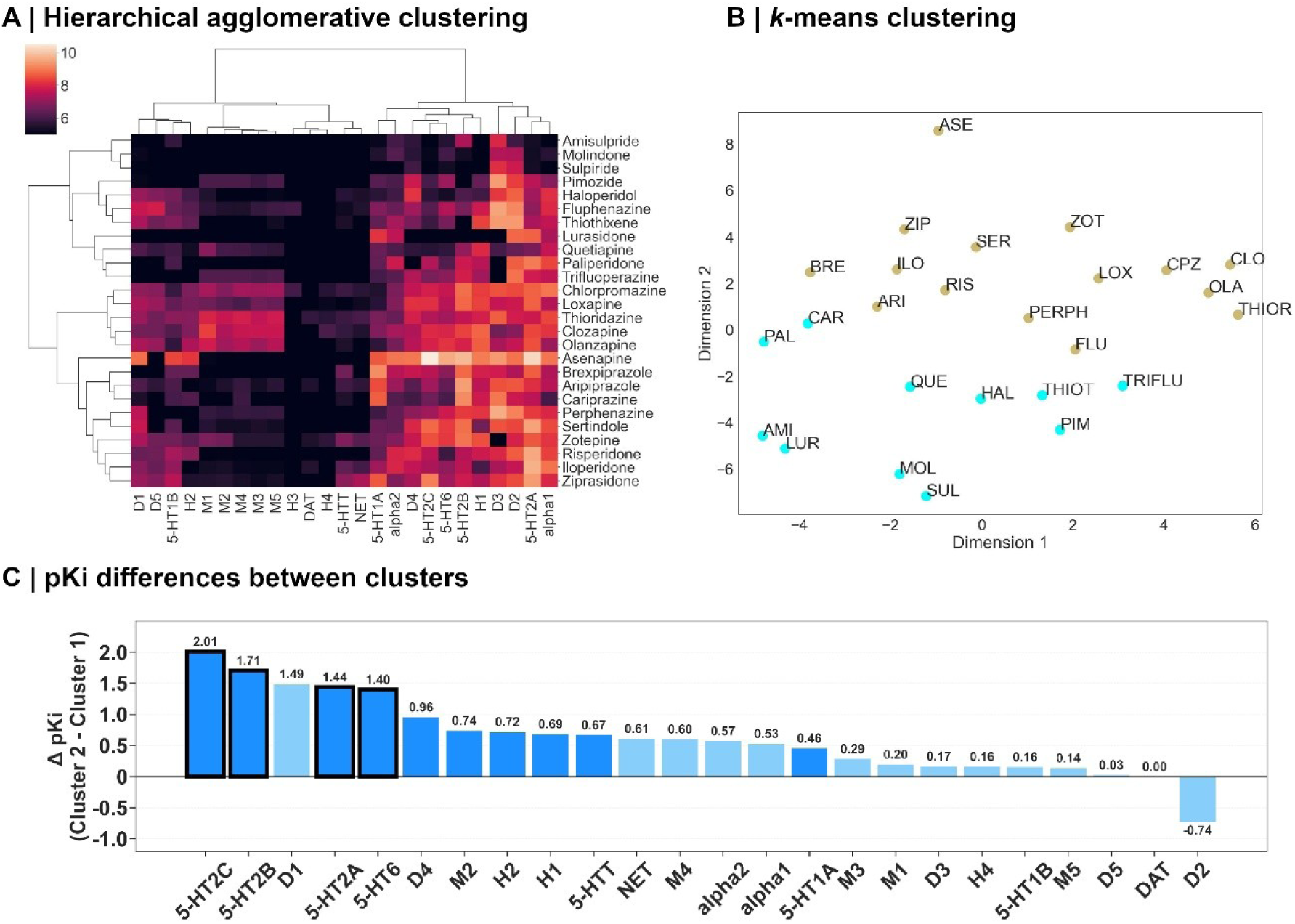
Influence of Clozapine and Quetiapine parent substances on clustering and between-cluster pKi differences. Replication of Figure 1 using the affinity profiles of the parent substances Clozapine and Quetiapine instead of the active metabolites Norquetiapine and Norclozapine.

## Bibliography

1. Jacobs R, Gutacker N, Mason A, Goddard M, Gravelle H, Kendrick T, et al. Determinants of hospital length of stay for people with serious mental illness in England and implications for payment systems: a regression analysis. BMC Health Serv Res. 2015;15:439.

2. Ye T, Huang W, Xu Z, Xu R, Yu P, Wu Y, et al. Greenness and hospital admissions for cause specific mental disorders: multicountry time series study. BMJ. 2025;391:e084618.

3. Barry SJ, Gaughan TM, Hunter R. Schizophrenia. BMJ Clin Evid. 2012;2012:1007.

4. Howes OD, Kapur S. The Dopamine Hypothesis of Schizophrenia: Version III—The Final Common Pathway. Schizophrenia Bulletin. 2009;35:549–562.

5. da Silva Alves F, Figee M, van Amelsvoort T, Veltman D, de Haan L. The revised dopamine hypothesis of schizophrenia: evidence from pharmacological MRI studies with atypical antipsychotic medication. Psychopharmacol Bull. 2008;41:121–132.

6. Osugo M, Whitehurst T, Erritzoe D, Carr R, Ashok AH, Maccioni L, et al. Role of Serotonin in the Neurobiology of Schizophrenia and Association With Negative Symptoms. JAMA Psychiatry. 2026;83:185–195.

7. Selvaraj S, Arnone D, Cappai A, Howes O. Alterations in the serotonin system in schizophrenia: a systematic review and meta-analysis of postmortem and molecular imaging studies. Neurosci Biobehav Rev. 2014;45:233–245.

8. de Jonge JC, Vinkers CH, Hulshoff Pol HE, Marsman A. GABAergic Mechanisms in Schizophrenia: Linking Postmortem and In Vivo Studies. Front Psychiatry. 2017;8:118.

9. Dalmau J, Armangué T, Planagumà J, Radosevic M, Mannara F, Leypoldt F, et al. An update on anti-NMDA receptor encephalitis for neurologists and psychiatrists: mechanisms and models. Lancet Neurol. 2019;18:1045–1057.

10. Hu W, MacDonald ML, Elswick DE, Sweet RA. The glutamate hypothesis of schizophrenia: evidence from human brain tissue studies. Ann N Y Acad Sci. 2015;1338:38–57.

11. Lahti AC, Weiler MA, Tamara Michaelidis BA, Parwani A, Tamminga CA. Effects of ketamine in normal and schizophrenic volunteers. Neuropsychopharmacology. 2001;25:455–467.

12. Seeman MV. History of the dopamine hypothesis of antipsychotic action. World J Psychiatry. 2021;11:355–364.

13. Kaul I, Sawchak S, Walling DP, Tamminga CA, Breier A, Zhu H, et al. Efficacy and Safety of Xanomeline-Trospium Chloride in Schizophrenia: A Randomized Clinical Trial. JAMA Psychiatry. 2024;81:749–756.

14. Laruelle M, Frankle WG, Narendran R, Kegeles LS, Abi-Dargham A. Mechanism of action of antipsychotic drugs: From dopamine D2 receptor antagonism to glutamate NMDA facilitation. Clinical Therapeutics. 2005;27:S16–S24.

15. Schwartz JC, Diaz J, Pilon C, Sokoloff P. Possible implications of the dopamine D(3) receptor in schizophrenia and in antipsychotic drug actions. Brain Res Brain Res Rev. 2000;31:277–287.

16. Velligan DI, Sajatovic M, Hatch A, Kramata P, Docherty JP. Why do psychiatric patients stop antipsychotic medication? A systematic review of reasons for nonadherence to medication in patients with serious mental illness. Patient Prefer Adherence. 2017;11:449–468.

17. Masand PS, Roca M, Turner MS, Kane JM. Partial Adherence to Antipsychotic Medication Impacts the Course of Illness in Patients With Schizophrenia: A Review. Prim Care Companion J Clin Psychiatry. 2009;11:147–154.

18. Goff DC. Maintenance treatment with long-acting injectable antipsychotics: comparing old with new. JAMA. 2014;311:1973–1974.

19. Blaisdell GD. Akathisia: a comprehensive review and treatment summary. Pharmacopsychiatry. 1994;27:139–146.

20. Wu H, Siafis S, Wang D, Burschinski A, Schneider-Thoma J, Priller J, et al. Antipsychotic-induced akathisia in adults with acute schizophrenia: A systematic review and dose-response meta-analysis. European Neuropsychopharmacology. 2023;72:40–49.

21. Huhn M, Nikolakopoulou A, Schneider-Thoma J, Krause M, Samara M, Peter N, et al. Comparative efficacy and tolerability of 32 oral antipsychotics for the acute treatment of adults with multi-episode schizophrenia: a systematic review and network meta-analysis. The Lancet. 2019;394:939–951.

22. Roth BL, Lopez E, Patel S, Kroeze WK. The Multiplicity of Serotonin Receptors: Uselessly Diverse Molecules or an Embarrassment of Riches? Neuroscientist. 2000;6:252–262.

23. Meltzer HY. What’s atypical about atypical antipsychotic drugs? Current Opinion in Pharmacology. 2004;4:53–57.

24. Zohar J, Stahl S, Moller H-J, Blier P, Kupfer D, Yamawaki S, et al. A review of the current nomenclature for psychotropic agents and an introduction to the Neuroscience-based Nomenclature. European Neuropsychopharmacology. 2015;25:2318–2325.

25. McCutcheon RA, Harrison PJ, Howes OD, McGuire PK, Taylor DM, Pillinger T. Data-Driven Taxonomy for Antipsychotic Medication: A New Classification System. Biol Psychiatry. 2023;94:561–568.

26. Creed-Carson M, Oraha A, Nobrega JN. Effects of 5-HT2A and 5-HT2C receptor antagonists on acute and chronic dyskinetic effects induced by haloperidol in rats. Behavioural Brain Research. 2011;219:273–279.

27. Laoutidis ZG, Luckhaus C. 5-HT2A receptor antagonists for the treatment of neuroleptic-induced akathisia: a systematic review and meta-analysis. Int J Neuropsychopharmacol. 2014;17:823–832.

28. Lieberman JA, Stroup TS, McEvoy JP, Swartz MS, Rosenheck RA, Perkins DO, et al. Effectiveness of Antipsychotic Drugs in Patients with Chronic Schizophrenia. New England Journal of Medicine. 2005;353:1209–1223.

29. Omdal R, Brokstad K, Waterloo K, Koldingsnes W, Jonsson R, Mellgren SI. Neuropsychiatric disturbances in SLE are associated with antibodies against NMDA receptors. Eur J Neurol. 2005;12:392–398.

30. Buck SA, Quincy Erickson-Oberg M, Logan RW, Freyberg Z. Relevance of Interactions Between Dopamine and Glutamate Neurotransmission in Schizophrenia. Mol Psychiatry. 2022;27:3583–3591.

31. Siafis S, Tzachanis D, Samara M, Papazisis G. Antipsychotic Drugs: From Receptor-binding Profiles to Metabolic Side Effects. Curr Neuropharmacol. 2018;16:1210–1223.

32. Berg KA, Clarke WP. Making Sense of Pharmacology: Inverse Agonism and Functional Selectivity. Int J Neuropsychopharmacol. 2018;21:962–977.

33. Khanna P, Suo T, Komossa K, Ma H, Rummel-Kluge C, El-Sayeh HG, et al. Aripiprazole versus other atypical antipsychotics for schizophrenia - Khanna, P - 2014 | Cochrane Library.

34. Wood MD, Scott C, Clarke K, Cato KJ, Patel N, Heath J, et al. Pharmacological profile of antipsychotics at monoamine receptors: atypicality beyond 5-HT2A receptor blockade. CNS Neurol Disord Drug Targets. 2006;5:445–452.

35. Peters L, Dyer M, Schroeder E, D’Souza MS. Invega Hafyera (Paliperidone Palmitate): Extended-Release Injectable Suspension for Patients With Schizophrenia. J Pharm Technol. 2023;39:88–94.

36. Jensen NH, Rodriguiz RM, Caron MG, Wetsel WC, Rothman RB, Roth BL. N-Desalkylquetiapine, a Potent Norepinephrine Reuptake Inhibitor and Partial 5-HT1A Agonist, as a Putative Mediator of Quetiapine’s Antidepressant Activity. Neuropsychopharmacol. 2008;33:2303–2312.

